# Identifying essential genes across eukaryotes by machine learning

**DOI:** 10.1101/2021.04.15.439934

**Authors:** Thomas Beder, Olufemi Aromolaran, Jürgen Dönitz, Sofia Tapanelli, Eunice O. Adedeji, Ezekiel Adebiyi, Gregor Bucher, Rainer Koenig

## Abstract

Identifying essential genes on a genome scale is resource intensive and has been performed for only a few eukaryotes. For less studied organisms essentiality might be predicted by gene homology. However, this approach cannot be applied to non-conserved genes. Additionally, divergent essentiality information is obtained from studying single cells or whole, multi-cellular organisms, and particularly when derived from human cell line screens and human population studies. We employed machine learning across six model eukaryotes and 60,381 genes, using 41,635 features derived from sequence, gene functions and network topology. Within a leave-one-organism-out cross-validation, the classifiers showed a high generalizability with an average accuracy close to 80% in the left-out species. As a case study, we applied the method to *Tribolium castaneum* and validated predictions experimentally yielding similar performance. Finally, using the classifier based on the studied model organisms enabled linking the essentiality information of human cell line screens and population studies.

## Introduction

Essential genes are defined as indispensable for the reproductive success of an organism (1). Consequently, information on essentiality is used in a broad range of life science research, prominently to identify drug targets, as in cancer therapy (2) or identifying insecticidal targets, but also for the design of a minimal genome in synthetic biology. Many genome wide screens exploring phenotypes and gene functions have been performed using forward genetic methods (3–5). Later, reverse genetic methods were developed which allowed targeting individual genes specifically (6). Some large scale screens were focused on the question, which genes are essential for an organism (7). Functionally, these screens revealed that essential genes are involved in fundamental cellular maintenance processes like DNA, RNA and protein synthesis (8). Besides this, their encoded proteins are highly connected in protein-protein interaction (PPI) and metabolic networks (9,10). Furthermore, core essential genes were identified across different model systems (11), revealing their evolutionary conservation. Many essential genes show their essentiality not as a binary trait nor is it fixed across all intrinsic and extrinsic conditions within the evolutionary niche and may be influenced by the environment and the genetic context (1). For example, in yeast many non-essential genes for growth in rich media are actually important in other growth conditions (12). Consequently, quantitative values for essentiality have been defined accounting for the degree of dependency on external influences, as well as the likelihood that a compensatory mutation occurs (13) and typically statistical score values are calculated for gene essentiality (14–16). Essential genes in humans were identified by studying cancer cell lines and more recently by genomic population studies, comprehensively reviewed by Bartha *et al*. (13). In cell lines, gene essentiality is assessed by cell viability after gene knock-out or knock-down, whereas in the population studies of humans it is assessed by scoring loss-of an allele or the depletion of variants in a gene. Interestingly, the essential gene lists of the two distinct approaches hardly overlap (13). This was unexpected since a cellular essential gene (CEG) should also be an organismal essential gene (OEG), even though not necessarily *vice versa*. This discrepancy between human CEG and OEG remains to be elucidated.

Despite their great value, experimental screens and population studies are very resource intensive. Consequently, on a genome scale, essential genes have been experimentally identified for several bacteria but only for few eukaryotes, while population studies were performed only for human (**Table 1**). Because of experimental challenges and costs, the computational prediction of essential genes is of great interest and machine learning can considerably facilitate the search for essential genes in an organism. Following this approach, classifiers have been trained on a set of genes with known essentiality that are described by various features. For this, features can be based directly on the DNA or protein sequence (17,18), such as GC content or amino acid frequencies, or on more complex characteristics e.g. the topology in PPI or co-expression networks (9,19). Subsequently, a trained classifier was used to predict a new set of genes finding novel essential gene candidates (9). The next milestone was the prediction of essential genes across species. For bacteria, the software Geptop 2.0 (20) calculates an essentiality score of an unknown gene based on information of 37 (bacterial) species and sequence similarity. For eukaryotes the task is more challenging, considering their complex multi-cellular architecture. Besides this, only few model organisms have been experimentally screened for essential genes to date, even though the number is growing (21,22). To our knowledge only two studies were published predicting essential genes across eukaryotes using machine learning. Lloyd *et al*. (23) predicted essential genes in two plant species and *S. cerevisiae*. They showed that inter-species prediction is feasible across plants, however, they observed a drastically reduced performance in cross plant-fungal species predictions. The other study, by Campos *et al*. (18), predicted essential genes in six eukaryotes using a leave-one-organism-out approach and it based merely on protein sequence features. These studies laid the foundations in machine learning based essential gene prediction highlighting the necessity to achieve a robust performance for predictions across organisms including humans. The aim of this study was to develop a classifier capable of identifying essential genes in eukaryotes even if no experimental data is available. For this, we trained and validated classifiers using essentiality information from six, well described model organisms. Within a leave-one-organism-out cross-validation, the classifiers were trained on data derived from five organisms and validated with the sixth organism. As a case study, we applied the classifier to the red flour beetle *T. castaneum*. We used the available RNA interference (RNAi) screen (24) with defined lethality status as a control, while validating further predictions experimentally. Moreover, we aimed to fill the gap between human cell line screens and population studies, by integrating information from the model organisms to improve human essential gene assignments.

**Table 1.** Assembled essential gene information based on OGEE (21), DEG (25) and the literature.

| Species | Source | Data set abbreviation | Method | Essential genes | Non essential genes |
| --- | --- | --- | --- | --- | --- |
| *H. sapiens* | Population sequencing data | Hs OE | Loss-of fuction-variants (33–37) | 2,828 | 12,844 |
| *H. sapiens* | Cell line | Hs CE | RNAi (26,30,47), shRNA (31), CRISPR (14–16) | 833 | 13,743 |
| *M. musculus* | Organism | Mm OE | Biallelic inactivation (39,46) | 1,966 | 4,505 |
| *M. musculus* | Cell line | Mm CE | CRISPR (40) | 939 | 7,472 |
| *D. melanogaster* | Organism | Dm OE | P-element (42), RNAi (27), CRISPR (41) | 246 | 271 |
| *D. melanogaster* | Cell line | Dm CE | RNAi (45), CRISPR (38) | 1,227 | 10,320 |
| *C. elegans* | Organism | Ce OE | RNAi (44) | 737 | 10,128 |
| *S. cerevisiae* | Single cell | Sc CE | PCR based (7,28,43) | 1,036 | 4,373 |
| *S. pombe* | Single cell | Sp CE | PCR based (29), Transposon (32) | 1,226 | 3,379 |
| Total |  |  |  | 11,038 | 67,035 |

## Materials and Methods

### Assembling the gold standard

We assembled essentiality information for genes from the six species *C. elegans, D. melanogaster, H. sapiens, M. musculus, S. cerevisiae* and *S. pombe*. For fly, mouse and human we could collect screens for CEG and OEG, for worm only for OEG, and for the yeasts (obviously) only for CEG. This essentiality information was collected from the databases OGEE (21) and DEG (25) and the literature as listed in **Table 1**. For genes with different essentially status in different screens, we performed a majority voting. For human cell line screens a gene had to be studied in at least five experiments. All genes, their class labels and predictions can be found in **Supplementary Table 4**.

### Feature generation

A total of 41,635 features were generated based on seven different sources including protein and gene sequence, functional domains, topological features, evolution/conservation, subcellular localization, and gene sets from Gene Ontology (**Figure 2**). Protein and gene sequences of the organisms were obtained using biomaRt (56). For genes with isoforms the features were generated individually for each isoform and the median of all was calculated. For deriving the protein and gene sequence features, various numerical representations characterizing the nucleotide and amino acid sequences and compositions of the query gene were calculated using seqinR (57), protr (58), CodonW (http://codonw.sourceforge.net/) and rDNAse (59). With seqinR simple protein sequence information including the number of residues, the percentage of physico-chemical classes and the theoretical isoelectric point were calculated. Most protein sequence features were obtained using protr including autocorrelation, CTD, conjoint triad, quasi-sequence order and pseudo amino acid composition. CodonW was used to calculate simple gene characteristics like length and GC content but also frequency of optimal codons and effective number of codons. With rDNAse DNA descriptors like auto covariance or pseudo nucleotide composition, and kmer frequencies (n=2-7) were calculated. Domain features including post-translational modifications were generated based on the tools provided by the Technical University of Denmark (http://www.cbs.dtu.dk/services/) and included prediction of membrane helices and β-turns, cofactor binding, acetylation and glycosylation sites. Topology features were computed based on PPAs derived from STRING v11 (60) including degree, degree distribution, betweenness, closeness and clustering coefficient using the Python library NetworkX. Conservation features included the number of homologous proteins of a query protein in the complete RefSeq (61) database using PSI-BLAST (62). The number of proteins found with e-value cutoffs from 1e-5 to 1e-100 (in 1e-5 multiplication steps) were used as features. In addition, an alignment coverage score (ACS) was calculated for hits with a cutoff ≤ 1e-30 as described by Vinayagam *et al*. (63). The ACS is the average of the query coverage score (QCS) and the subject coverage score (SCS):

QCS and SCS are combined measures of size, identity and E-value of the alignment concerning the query or subject sequence,

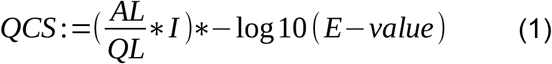

and

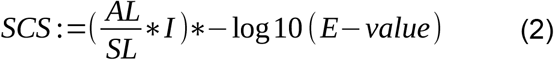

where AL denotes the alignment length, QL the length of the query sequence, SL the length of the subject sequence, and I the fraction of identical amino acids in the alignment. Next the number of homologous sequences with a score from 0 to 0.95 in 0.05 steps were calculated. Identically, the number of paralogous sequences were calculated, but blastn (64) alignment results with an e-value cutoff ≤ 1e-30 were used as input for the score. Subcellular localization features were predicted using DeepLoc (65), which assigns a score for each protein to its localization in eleven eukaryotic cell compartments. Gene set features were computed based on 3,874 Gene Ontology (GO) terms present in all analyzed organisms similar to Chen el al. (52). By this, not only the characterization of the query gene was taken into account, but also of its neighbors in the protein association network making the features more robust against false gene set annotations. We assembled the neighbors of the query gene employing the gene network definitions STRING v11 (60). For each of the gene sets, a Fisher’s exact test for enrichment of interaction partners was performed. The -log10 values of the p-values were used as features.

### Data normalization, feature selection and machine learning

Data analysis was performed using R. Values of each feature were z-transformed and each value was assigned to deciles. Next, we performed two steps for feature selection prior to the training procedure. The data was randomly split into training (4/5) and testing (1/5). Based on the training set, Least Absolute Shrinkage and Selection Operator (LASSO) was applied for feature selection using the glmnet package (66) in R (cv.glmnet function with parameters alpha = 1, type.measure = “auc”). To avoid collinearity, highly correlating features with Pearson correlation coefficients r ≥ 0.70 were removed.

To overcome class imbalances when training the classifiers, we used SMOTE (67) and trained with the classification algorithms Random Forest (RF) and Extreme Gradient Boosting (XGB) from the caret (68) package. For RF the *tuneLength* parameter in the *train* function was set to 3 resulting in 3 *mtry* values (number of predictors randomly sampled at each split). For XGB *eta, nrounds, max_depth*, min*_child_weight* and *colsample_bytree* were optimized in a tune-grid whereas *gamma* and *subsample* parameters were kept constant at 0 and 1, respectively. This resulted in 216 different parameter combinations for XGB tuning. To improve generalizability, we performed a stratified randomized 5-fold cross-validation. Thereby, 80% of the data was used for feature selection, hyperparameter tuning and training of the classifiers, and 20% for testing (**Supplementary Figure 1**). Within the training step, features were selected, the model was learned and evaluated in an inner 5-fold cross-validation (inner loop).

### Leave-one-organism-out cross-validation scheme for the CLassifier of Essentiality AcRoss EukaRyotes (CLEARER)

For each individual species (five species for CEG, four for OEG), five machines were trained (**Supplementary Figure 1**). The left-out species were predicted with machines trained on the according CEG or OEG data sets of the other organisms. Thereby the classifiers for each (non-left out) species gives an essentiality prediction score between zero and one and the average of these scores was used for the prediction of a gene to be essential in the left-out species.

### Orthology-based essential gene prediction

We derived orthologs from the OrthoDB v10 (69) database and assigned the essentiality to the genes according to the data sources listed in **Table 1**. For predicting essential genes in an organism we selected the essential and non-essential orthologs in the other organisms and performed a hypergeometric test. P-values were FDR corrected for multiple testing and values lower than 0.05 considered to be significant.

### RNAi experiments in *T. castaneum*

RNAi was performed according to the procedure described for the larval injection screen in Schmitt-Engel *et al*. (24) with minor modifications: We used another strain (San Bernardino) and scored lethality after 7 days after injection (instead of 11). We defined a gene as lethal if the lethality in the pupal or larval screen was at least 50%.

### Functional enrichment analysis of essential genes

To study in which cellular processes essential genes were enriched, an enrichment analysis was performed using Gene Ontology version from 2020-11-18, biological process, molecular function and cellular compartment. Enriched GO-terms (Fisher’s exact test p < 0.05, FDR corrected) were selected and compared across the six species. Gene sets were removed if they showed high redundancy according to the following method. Redundancy between two gene sets was quantified using Jaccard similarity coefficients,

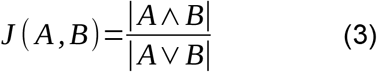

in which A and B are gene sets enriched for essential genes. An undirected graph G = (X,E) is introduced, with X being gene sets as vertices and E being gene set pairs with J(A,B) >= 0.3 as edges of the graph. A mixed integer linear model (weighted stable set problem) was setup with a constraint for each edge to select at most one of the vertices of an edge:

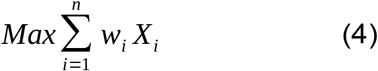

subjected to

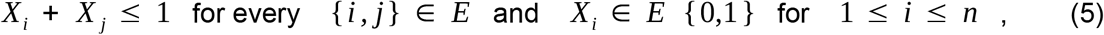

where, w_i_ is the weight of a gene set. The weight is derived from the enrichment test p-value and maximization was performed employing linear integer programming solved by the software Gurobi (version7.5.2https://www.gurobi.com). This led to an optimal selection of at most one gene set from a pair in such a way that the overall number of non-redundant gene sets were maximized. Moreover, very general gene sets containing ≥ 1000 genes (in any organism) were removed and gene sets comprising ≤ 0.1% genes of the corresponding organism were not considered. For illustration, each gene set was assigned to one of eleven major groups (cell cycle, cellular structure, development, immune response, metabolism, neural processes, protein biogenesis, regulation, repair, RNA biogenesis and signaling).

### Clustering

Scores of human cell lines were combined calculating the rank products leading to a combined rank for CEG. Similarly, combined ranks were calculated for OEG based on the scores of the population studies. Clustering of human genes was based on the percentiles of the combined scores from the cell lines (experimental CEG), population studies (experimental OEG) and CLEARER (OEG and CEG predictions). Clustering was performed using euclidian distance and average linkage of the R package pheatmap.

### Associating human genes to phenotypes

To investigate how the predicted essential genes in human associate with human diseases, we extracted 225,443 phenotype-to-gene associations from the Human Phenotype Ontology (53) database. For each of the phenotypes an enrichment test (Fisher’s Exact Test) for the according genes in the clusters was performed following FDR correction and phenotypes with p < 0.05 were considered to be significantly enriched. Word clouds illustrating phenotypes were generated using the R package wordcloud2.

## Results

### Essential genes are different in single cells compared to multi-cellular organisms

Our study was performed based on the six model organisms *Homo sapiens* (human), *Mus musculus* (mouse), *Drosophila melanogaster* (fly), *Caenorhabditis elegans* (worm), *Saccharomyces cerevisiae* and *Schizosaccharomyces pombe* (yeasts). Essentiality information for each of these organisms was taken from the databases Online GEne Essentiality (21), Database of Essential Genes (25) and the literature (7, 14–16, 26–47). **Table 1** lists the number of genes for which this essentiality information could be assembled. To our knowledge we compiled the most comprehensive collection of essentiality information for these six eukaryotes comprising 11,038 essential and 67,035 non-essential labeled genes.

For humans, it has been shown that essentiality information derived from viability screens of cancer cell lines does not well agree with essentiality information derived from *in vivo* genetic studies (13). We speculated that this phenomenon can be generalized, i.e. CEG identified from cell lines or unicellular organisms (yeasts) are distinct from OEG. E.g. genes involved in embryo development or neural morphogenesis may be substantial for multicellular organisms, but not for a unicellular organism or a cell line. Hence, we defined two categories of gene essentiality i.e., CEG and OEG, depending on the cellular or organismal nature of the experimental study (**Table 1**). We compared gene essentiality based on OEG and CEG experiments across the six investigated organisms based on orthologous gene groups. As shown before (13), gene essentiality inferred from human *in vivo* population studies correlated well with each other (r = 0.53 ± 0.15) and the correlation was even better between human cell line studies (r = 0.74 ± 0.07, **Fig. 1**). Interestingly, we also observed good pairwise correlations of CEG of human, fly, mouse and the yeasts (r = 0.38 ± 0.17). Moreover, we observed reasonable pairwise correlations of OEG across organisms (r = 0.20 ± 0.06). In contrast, there was only low correlation between CEG and OEG across organisms (r = 0.15 ± 0.13) and the lowest correlation between OEG and CEG of the same species was found in human (r = 0.13 ± 0.06). The complete list of orthologous groups and essentiality information is provided in **Supplementary Table 1**.

**Fig. 1.**
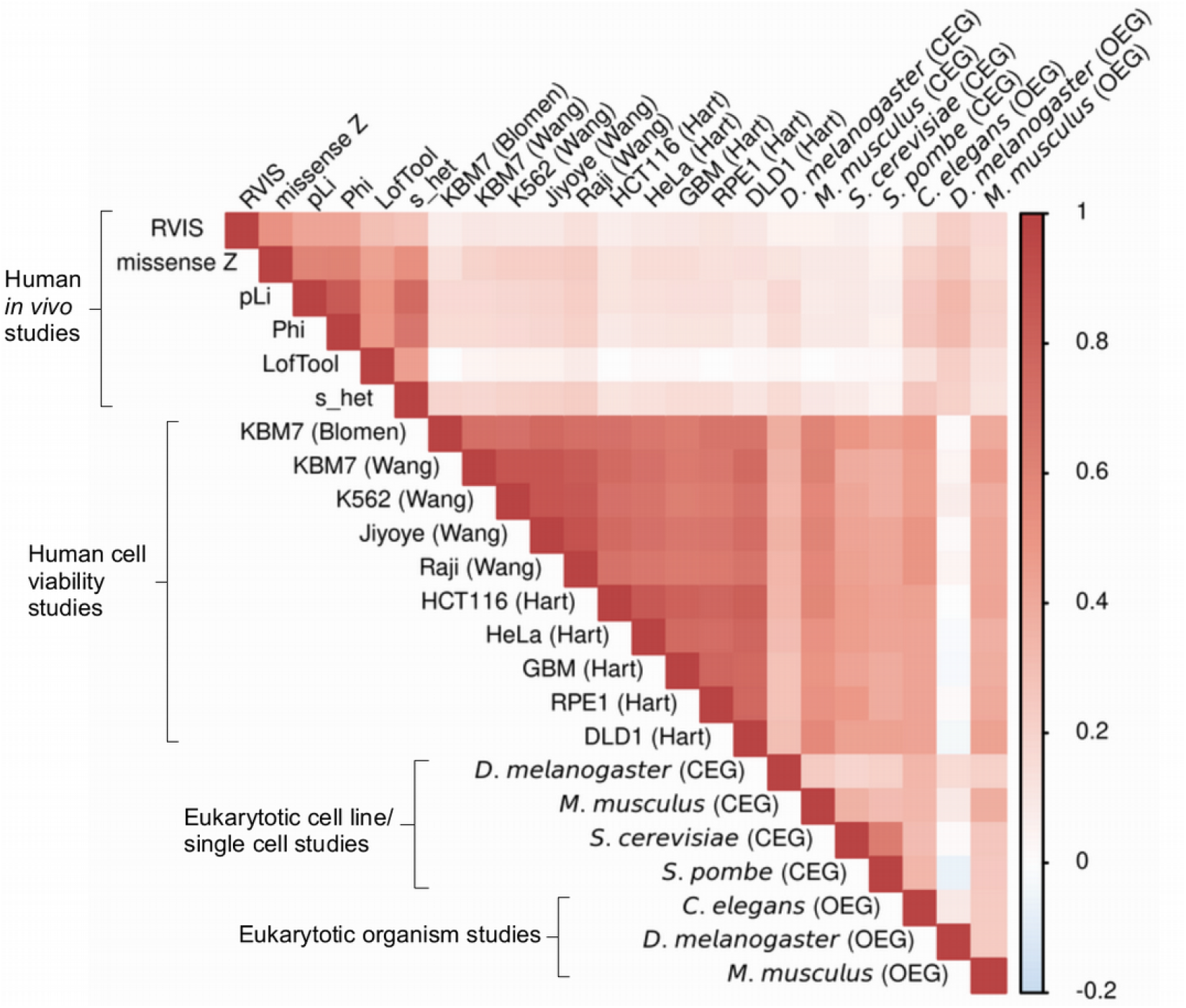
Correlation of cellular and organismal essential genes. Correlation analysis of essentiality across organisms shows that both cellular (CEG) and organismal essential genes (OEG) are conserved across the investigated eukaryotes. Essentiality on the cellular level correlated better than on the organismal level. The lowest correlation of CEG and OEG was observed for human. Human OEG scores (from population studies) and human CEG scores (from cell based knockout/knockdown screens) were obtained from Bartha *et al*. (13). Human population studies are denoted as scores used in to define essentiality. The human cell line studies are denoted by the name of the corresponding cell lines and, in brackets, the first author of the study. Additionally, the OEG and CEG scores for *C. elegans, D. melanogaster, M. musculus, S. cerevisiae* and *S. pombe* were included.

Next, we studied the involvement in cellular processes of CEG and OEG. For this, we performed gene set enrichment analyses based on the gene set definitions of Gene Ontology. To get a better overview, each gene set was assigned to one of eleven major groups (cell cycle, cellular structure, development, immune response, metabolism, neural processes, protein biogenesis, regulation, repair, RNA biogenesis and signaling).

Whereas CEG were enriched in processes describing cellular macromolecule biogenesis and cell cycle/proliferation, OEG showed enrichment in regulation, development/morphogenesis, neural related processes and signaling (**Fig. 2**). This reflects the need of multicellular organisms for functional organ systems, which do not only depend on the survival of the respective cells but also their concerted function within and between organs. **Fig. 2** shows the proportions of CEG and OEG of these different cellular processes for each organism. Notably, for human OEGs less developmental gene sets were found compared to *M. musculus, D. melanogaster*, and *C. elegans* potentially reflecting the different way to identify essential genes. The simple multicellular nematode *C. elegans* shows processes we observed in CEG of the other multi cellular organisms. All enriched gene sets are listed in **Supplementary Table 2**.

**Fig. 2.**
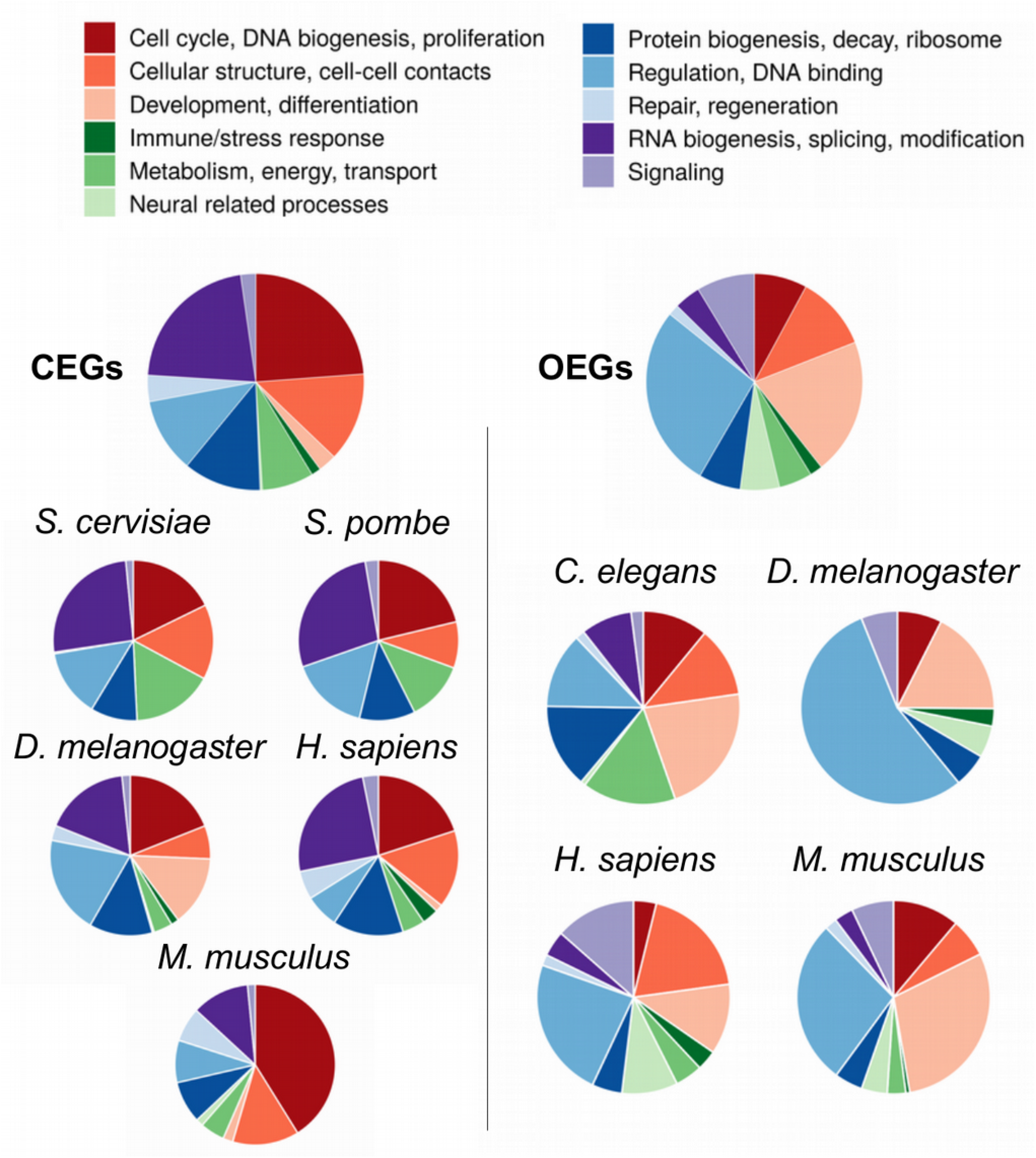
Functional characterization of essential genes. Distribution of CEG and OEG in major biological processes. Enriched gene sets were assigned to one of eleven major categories. The proportions were derived by dividing the number of essential genes in each gene set by the total number of essential genes of the according CEG or OEG entity. CEG were enriched in processes describing cellular biogenesis and cell cycle/proliferation, whereas OEG showed enrichment in regulation, development/morphogenesis and signaling.

We observed that CEG are similar among the individual species, and similarly OEG. However, CEG and OEG are substantially different suggesting learning our machines with CEG and OEG separately.

### Setting up the data sets for machine learning

We set up a machine learning procedure to predict essential genes across eukaryotes called **CL**assifier of **E**ssentiality **A**c**R**oss **E**uka**R**yotes (CLEARER). Each data set from **Table 1** served as the gold standard, and with this we trained individual classifiers for each organism for CEG and OEG based on 41,635 features from seven categories comprising information from protein and DNA sequences, domains, of gene network topology, evolutionary conservation, subcellular localization, biological processes and further gene set definitions (**Fig. 3**). The complete list and description of features used in this study is shown in **Supplementary Table 3**. To predict essential genes across species, a leave-one-organism-out cross-validation was applied. An example is shown in **Fig. 3** where CEG of human are predicted by machines trained on fly, mouse and yeasts. The machine learning workflow is depicted in **Supplementary Figure 1**. Notably, following a leave-one-organism-out cross-validation allowed realistic performance evaluations as the class labels of the left-out organism were only uncovered when evaluating the predictions. OEG were predicted separately, using essentiality information of the four available organisms comprising worm, fly, mouse and human (**Table 1**).

**Fig. 3.**
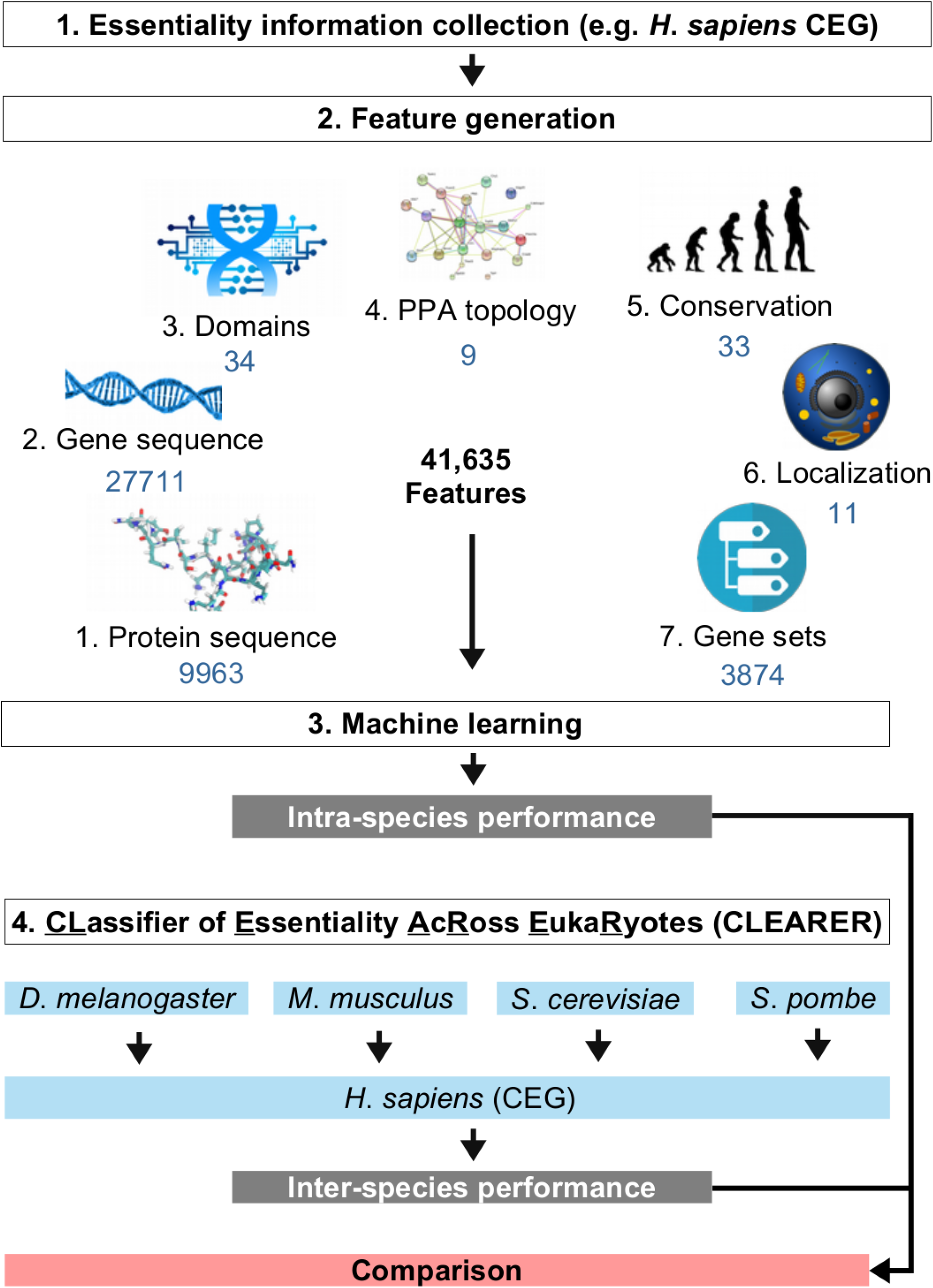
The workflow of CLEARER. The workflow is exemplified for the prediction of human cellular essential genes (CEG). For each gene 41,635 features based on seven different categories were assessed. Machine learning was performed and the intra-species classification performance evaluated. For the inter-species classification, the machines trained on the other organisms were used to predict human CEG.

### Identifying essential genes within and across species

First, we tested our approach by predicting essential genes within the same species using a stratified randomized five-fold cross-validation. Thereby, 80% of the data was used for feature selection, hyperparameter tuning and training of the classifiers, and 20% for testing (**Supplementary Figure 1**). We evaluated the performance of the two machine learning strategies Random Forests and Extreme Gradient Boosting. Random Forests performed slightly better on the test sets (**Supplementary Figure 2**) with an average ROC-AUC of 0.857 ± 0.057. The best performance was obtained for the human cell lines (ROC-AUC = 0.955 ± 0.018), which also reflects the consistency of the essentiality information within the six data sets (**Table 1**). Notably, we observed no difference between the performance in predicting CEG and OEG. For both, we got consistently high performances (**Fig. 4a**, ROC-AUC_CEG_ = 0.873, ROC-AUC_OEG_ = 0.845, p = 0.13).

**Fig. 4.**
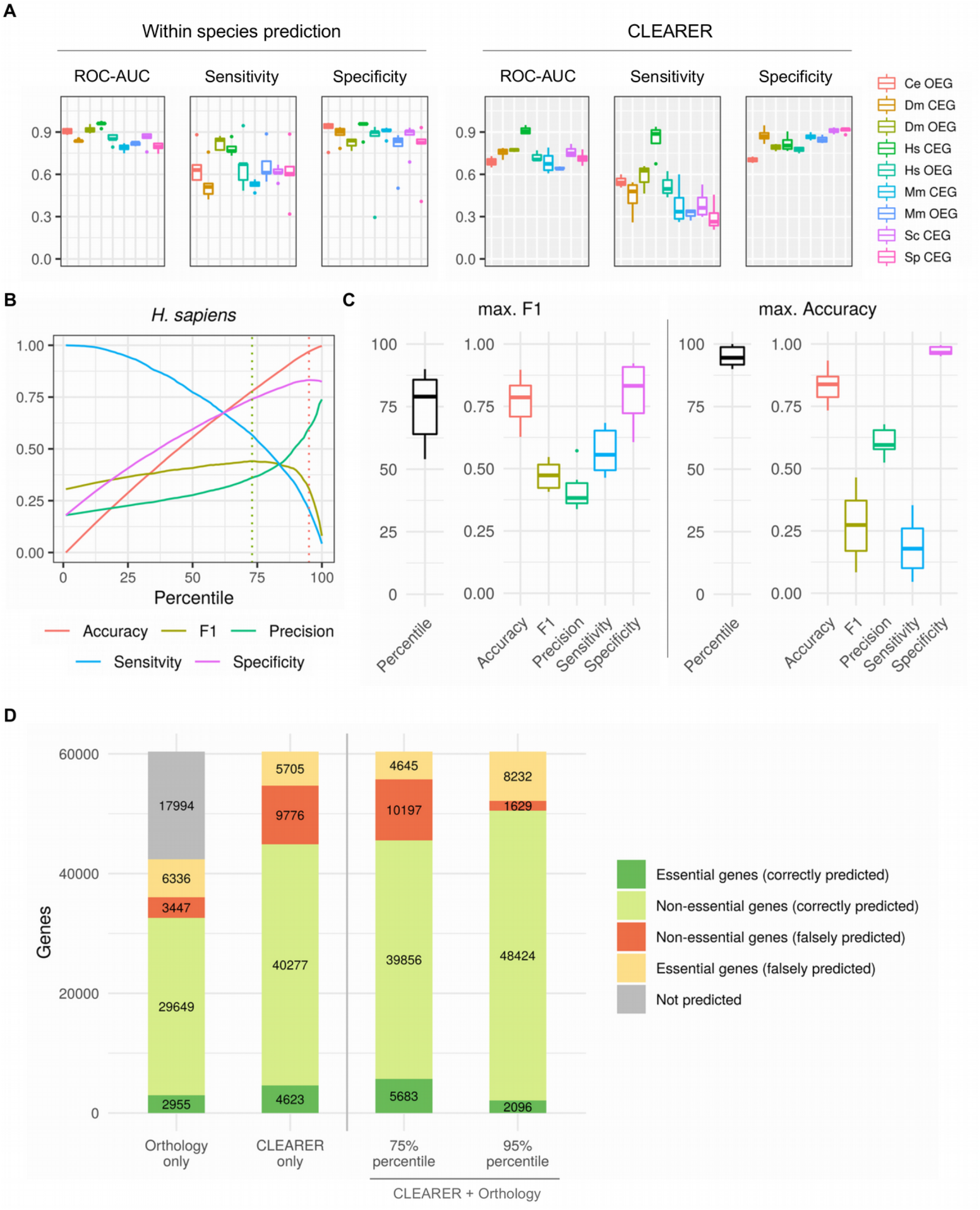
Performance of CLEARER. **A** Comparison of the prediction performance within and across species from all cross-validations. Species abbreviations are as listed in **Table 1. B** Line graph illustrating the maximal F1-score and accuracy cutoff for *H. sapiens* after combining CLEARER and the orthology-based approach. Dotted lines indicate maxima. **C** Box plots showing the percentiles and performance metrics for the maximal F1-score and maximal accuracy for the six model organisms. **D** Bar graphs with the total number of correct and incorrect predicted genes.

Furthermore, we investigated the performance when combining CEG and OEG information. For this, we trained and tested using combined lists. As expected, we observed a distinctively reduced performance in terms of the ROC-AUC for the organisms with CEG and OEG information (**Supplementary Figure 3**). For the following, we based our analysis on classifiers which were trained separately either with OEG or CEG. In conclusion, our method predicted essentiality well, when CEG and OEG were trained separately and were applied to the same species. Next, CLEARER (**Fig. 3**) was applied to predict CEG and OEG across organisms, again following a leave-one-organism-out cross-validation. We observed an ROC-AUC of 0.744 ± 0.084 on average (**Fig. 4a, Supplementary Figure 4** shows the performance for each organism separately), and the overall accuracy of the final predictions of the within species classifiers and CLEARER were similar (no significant difference). Next, we compared CLEARER to the approach previously reported by Campos *et al*. (18), which used protein sequence features and the leave-one-organism-out cross-validation approach investigating the same species. In their approach CEG and OEG were not handled separately and machines were not trained on the individual organisms, but on combined lists of essential and non-essential genes (18). Following this way, we observed a reduced performance compared to CLEARER when using the same protein sequence features (ROC-AUC = 0.599 ± 0.067, p < 0.0001) and also when using our set of diverse features (ROC-AUC = 0.696 ± 0.044, p < 0.05). This demonstrated the improvement when integrating the comprehensive set of the above described features and an ensemble classifier trained on CEG and OEG of the individual species.

CLEARER performed well to predict essential genes for each of the model organisms investigated when trained on the other model organisms and significantly outperformed the previous approach.

### Comparing CLEARER with an orthology-based prediction approach

The most common approach to find essential genes is based on inference by orthology. For all genes with orthologs in at least one of the other investigated organisms we performed a test for enrichment of essential orthologs. This enrichment test showed a similar accuracy as simple majority voting (accuracy_enrichment_ = 0.784 ± 0.052, accuracy_majority_ = 0.797 ± 0.047, p = 0.1563) but an increased sensitivity (sensitivity_enrichment_ = 0.398 ± 0.188, sensitivity_majority_ = 0.314 ± 0.141, p = 0.031). Consequently, we used the enrichment test-based assignment. A major disadvantage of such an approach is that for 17,994 genes (29.8%) no ortholog with known essentiality was assigned, leaving these genes unpredicted (**Fig. 4d**). However, for the genes with identified orthologs the enrichment test performed similar to CLEARER in terms of the accuracy (accucacy_enrichment_ = 0.77, accucacy_CLEARER_ = 0.74, **Fig. 4d**). Despite that, CLEARER classified more essential (31.6%) and non-essential genes (24,4%) correctly (**Fig. 4d**). Notably, both approaches shared a substantial number of true positive predictions (**Supplementary Figure 5**) suggesting combining both approaches for the best performance.

### Combining CLEARER with the orthology-based approach improves the overall predictions

To obtain a unique classifier predicting CEG and OEG with the best performance, we now combined the predictions from CEG, OEG and the orthology-based approach. For the yeasts only predictions for CEG and from the orthology-based approach were considered. A combined score for each gene was calculated based on the ranking of the predictions for each approach. This score allowed to order the genes such that an optimal percentile (cutoff) could be selected above which a gene was regarded to be essential. The optimal cutoff needed to be selected to balance between high accuracy, sensitivity and precision. For this, two measures were regarded, i.e. (1) the maxima of the F1-score (harmonic mean of sensitivity and precision) and (2) the maximal accuracy. **Fig. 4b** illustrates this tradeoff exemplarily for the prediction of essential genes for human. Across all six organisms, the best cutoff based on the maximal F1 approach was on average the 75% percentile, yielding an accuracy of 0.769 ± 0.99 (**Fig. 4c**). The best cutoff for the maximal accuracy was the 95% percentile and yielded a higher accuracy (0.832 ± 0.077) and precision (0.606 ± 0.059), but the sensitivity was lower (0.186 ± 0.117). Even though the latter criterion yielded lower sensitivities, depending on the application, high precision can be very beneficial if validation experiments are costly or technically complex. We observed improved predictions (compared to CLEARER alone) with the combined approach yielding n = 1,060 (18.7%) more correctly identified essential genes and only a marginal decrease in specificity (1.1%) when applying the 75% percentile cutoff (**Fig. 3d**). We used this combined classifier to predict essential genes for the case study application (next section). The results of the predictions for all model organisms are shown in **Supplementary Table 4**.

### CLEARER combined with the orthology-based approach performs well for *Tribolium castaneum*

As a case study, we applied the unified classifier (CEG_CLERARER_, OEG_CLEARER_ and orthology-based approach) to *T. castaneum. T. castaneum* is an insect with emerging interest because in many respects its biology is more representative of insects than that of *D. melanogaster* (48), e.g. *T. castaneum* is used as a model organism for pest control (49). Gene essentiality information for is available from previously published knockdown screens (24, 49, 50), and was used to validate the predictions. We made predictions for 12,859 genes of which 2,783 had been tested by RNAi (**Supplementary Table 5**). Following the F1-based cutoff criterion, the top 25% predictions were considered to be essential. In total 610, essential genes were correctly identified with a precision of 0.712 (specificity = 0.809), showing that over 70% of predicted essential genes are indeed essential. The overlap of essential gene predictions and experimental results was highly significant (p < 0.0001, odds ratio = 2.63). Following the maximal accuracy cutoff, the top 5% of predictions were considered to be essential yielding high precision (0.859) and specificity (0.973), but only low sensitivity (0.143). The results show that CLEARER predicts essential genes for *T. castaneum* with a very similar performance as achieved for the model organisms when applying the leave-one-organism-out cross-validation. Next, we selected 200 genes with the highest prediction scores and validated them experimentally performing RNAi *in vivo*. Indeed, n=160 genes (accuracy = 80.5%) proved experimentally to be essential (**Supplementary Table 5**). In addition, we randomly selected 200 genes and tested their essentiality *in vivo* (**Supplementary Table 5**). Again, the overlap of essential gene predictions and RNAi validation experiments was significant (p < 0.01, odds ratio = 2.78, precision = 0.646 and specificity = 0.850).

In summary, applying our approach to *T. castaneum* yielded a similar performance as observed in the leave-one-organism-out cross-validation of the six model organisms showing the good applicability of our approach.

### CLEARER supports defining human essential genes

Essential gene information for human derived from cancer cell line and population sequence studies hardly overlap, indicating a missing link between both approaches. We aimed to provide this link by integrating the available experimental information for human with predictions from the model organisms.

Essentiality scores of the experimental data from ten human cell line screens were combined (using rank products) to obtain an experimental CEG score. Similarly, a combined score was obtained for OEG based on the five population studies. As described above, human CEG and OEG poorly overlap, and hence the correlation of the according scores was low (r = 0.12, **Fig. 5a**). In contrast, the scores from the combination of CLEARER and the orthology-based approach correlated much better with the scores from the cell line screens (r = 0.50) and the population studies (r = 0.24). Next, experimentally derived CEG and OEG scores were combined with the computational predictions using the rank product. This combined score showed the best correlation to the cell line screens (r = 0.53), population studies (r = 0.45) and CLEARER (r = 0.71).

**Fig. 5.**
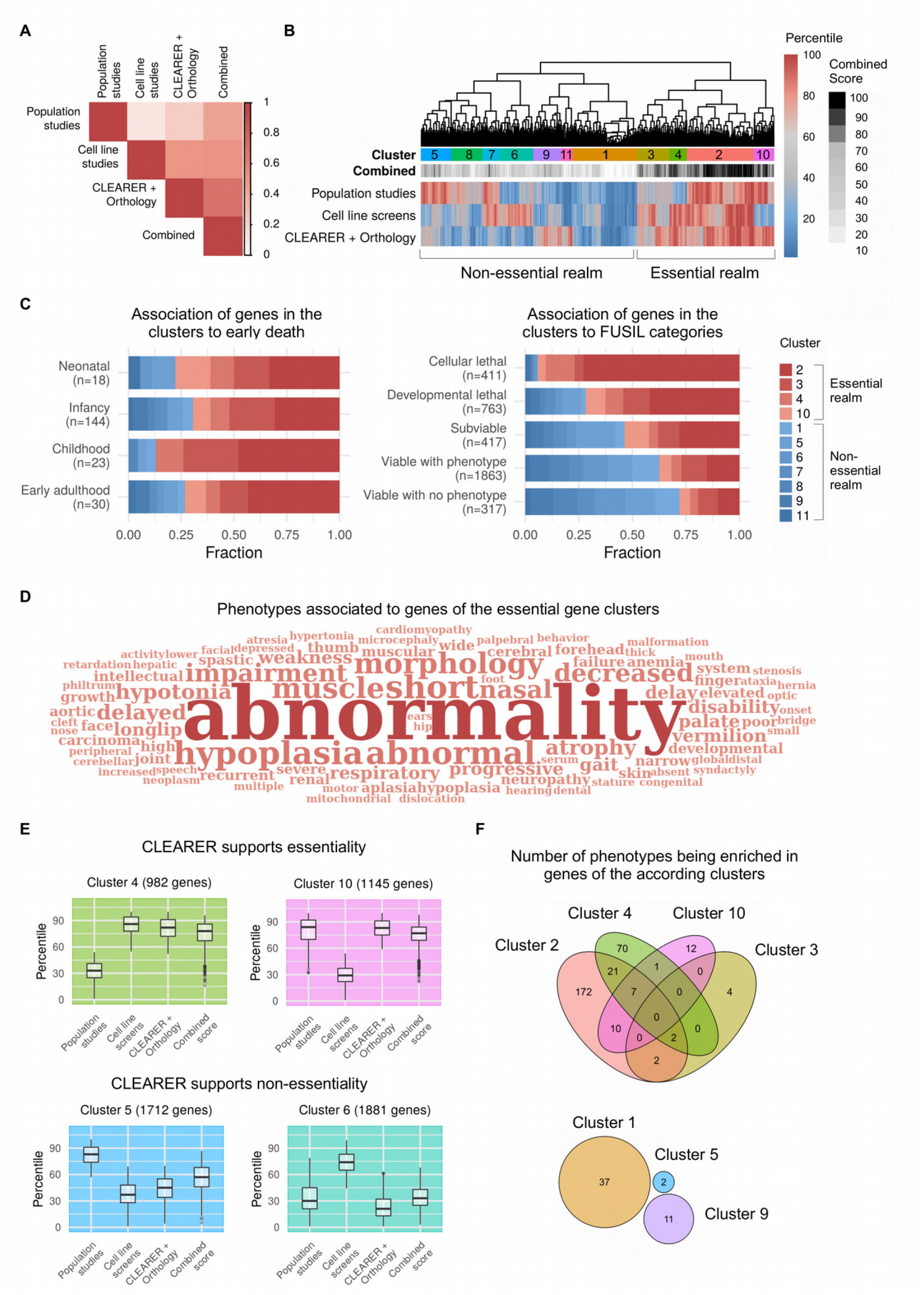
CLEARER supports identifying essential genes for human. **A** Correlation of experimental and computational scores. Scores from five population studies and ten cell line screens were combined and compared to predictions from CLEARER. The highest correlation was found when combining the experimental data and computational predictions. **B** Clustering of computational and experimental scores separates essential and non-essential genes. **C** Essential gene clusters associate with early death and lethality from Cacheiro *et al*. Full Spectrum of Intolerance to Loss-of-function categories (51). **D** Word cloud illustrating enriched phenotype-to-gene associations of genes from the essential gene clusters. **E** Box plots of the scores of the four essential gene clusters illustrate how CLEARER supported the final decision-making towards or against essentiality. **F** Venn diagrams showing the number of overlapping and non-overlapping phenotypes being enriched in genes of the according clusters. Notably, for the non-essential gene clusters 6,7,8 and 11 no enriched phenotype was identified.

We were now interested whether this combined score improved elucidating human essential gene definitions. For this, we clustered the essential genes based on their experimentally derived scores and CLEARER predictions. We identified eleven clusters distinctively separating essential and non-essential genes according to their combined scores (**Fig. 5b**). In total, 7,739 genes (38.9%) with high combined essentiality scores were found in four clusters (2, 3, 4 and 10). Genes from these four clusters account for the majority of genes associated to an early death (72.6%, **Fig. 5c**). Cacheiro *et al*. (51), recently proposed a categorization of human genes according to their essentiality. Genes from cluster 2, 3, 4 and 10 were highly enriched in the categories “cellular lethal” and “developmental lethal” of Cacheiro *et al*. (**Fig. 5c**). In contrast, these genes were depleted in the “viable” categories, which again indicates an accumulation of essential genes in cluster 2, 3, 4 and 10. Another indicator for essentiality was their association to human diseases. Genes from our essential gene clusters were combined and compared to genes from the non-essential clusters (clusters 1, 5, 6, 7, 8, 9 and 11). We observed a distinctively high enrichment of diseases and impairments associated to the genes from essential gene clusters. In total 490 phenotypic descriptions were found to be enriched mostly associated with abnormal development but also hypoplasia, which is associated with an inadequate or below-normal number of cells (**Fig. 5d**). In contrast, the 12,175 genes from the other seven (non-essential gene) clusters were not enriched for any phentoypic description. These results highly suggest a strong enrichment of essential genes in clusters 2,3,4 and 10, compared to the other clusters.

Another interesting finding was that in clusters where experimental and computational scores differed, CLEARER aided in making the final decision. Genes from cluster 4 and cluster 10 had high scores either in the cell line screens (cluster 4) or population studies (cluster 10). Both were supported by the predictions from CLEARER. In contrast, genes from cluster 5 and 6 were not supported by CLEARER (**Fig. 5e**). Next, the phenotypic associations of the individual clusters were compared (**Fig. 5f**) again showing the overlap of associated diseases in the essential gene clusters. On the contrary, genes from the non-essential clusters were considerably less associated to diseases (contributing only 3.6% of all identified phenotypes) and their phenotypes didn’t overlap (**Fig. 5f**). In fact, for the non-essential gene clusters 6, 7, 8 and 11 no associated phenotype was found. The combined score for each gene and enriched phenotypes for the clusters are listed in **Supplementary Table 6**.

The results show that the combined score supports defining human essential genes and may contribute filling the gap between cell line screens and population studies by integrating information from the other model organisms. Genes with high scores are associated with death, abnormal morphology, cancer and other diseases. Moreover, the combined score appears to be particularly supportive when the results of the cell line screens and population studies are divergent.

## Discussion

For prokaryotes, prediction of essential genes is possible with a tool that uses sequence comparisons (20). Essential gene prediction for eukaryotes is more challenging as less genome wide experimental screens are available and the approaches are heterogeneous. Experimentally, they comprise knock-out as well as knock-down methods investigating cell lines, single cell organisms, whole multi-cellular organisms and population studies.

Consequently, this leads to rather divers lists of identified essential genes. To cope with these inconsistencies we followed two principles. Firstly, we consolidated various studies (26 in total) for the individual organisms thus balancing for differences. Secondly, predictions were based on machines trained across several organisms. Consequently, several methods and biological settings were considered integrating these frameworks. In line, predictions with about 80% accuracy were consistently achieved across the six studied model organisms. Our approach highly outperformed a previously published method (18), by employing a broad range of gene descriptors derived from gene and protein sequences, protein domains, evolutionary conservation estimated from homology analyses, cellular processes and functions, but also from topology descriptors of gene association based networks. Moreover, we showed the advantage of distinguishing between CEG and OEG.

Particularly for human, there is a large discrepancy between the available essentiality from cell line screens compared to population studies (13). We showed that combining essentiality information from our machine learning pipeline CLEARER, orthology and experimental data for human led to a combined score highly correlating with scores from both, cell line and population studies. Genes with high scores associated with early death, abnormal morphology and cancer. The predictions from CLEARER supported the final decision of genes with higher scores from the cell line or population studies. This suggests that the combined score, integrating experimental data and computational predictions based on the model organisms, provides an important resource for genetic studies of human health and diseases.

Recently, Cacheiro *et al*. (51), binned essential gene information of mouse and human orthologous genes resulting in 3,819 predicted human essential genes. The assignment of their groups is in accordance to our combined score. However, the clear advantage of our approach is that it is applied on a genome scale and not just to orthologs. HEGIAP (52) a collection of analytical tools enabling to analyze the epigenectics, gene structure and evolution of human essential genes was published recently. However, HEGIAP considers only human essential genes from cell line screens neglecting organismal essentiality. Besides this, HEGIAP is rather a descriptive tool collection not providing a conclusive prediction for gene essentiality. Our method allows the prediction of essential genes, in principle, for any eukaryote without the need of a screening experiment. This can simplify the search for essential genes in non-model organisms e.g. to find targets for pest and vector control. Instead of large scale experimental screens, CLEARER can provide a shortlist of putative essential genes, which can subsequently be tested experimentally on a smaller scale.

In this study we focused solely on the prediction of essential genes. As a perspective, using the same approach, also other gene to phenotype associations can be predicted. Several databases with gene to phenotype descriptions for model organisms exist (39, 50, 53–55) and can be used for setting up the gold standard. Moreover, here we only considered loss-of-function of single genes. Predicting synthetic lethality, in which combinations of loss-of-functions lead to death can be a further promising application.

## Data Availability

Source code is available at https://github.com/ThomasBeder/CLEARER.

## Funding

This work was supported by the Deutsche Forschungsgemeinschaft (https://www.dfg.de/) within the project KO 3678/5-1 and the German Federal Ministry of Education and Research (BMBF) within the project Center for Sepsis Control and Care (CSCC, 01EO1002 and 01EO1502). Additionally, the work was supported by Bayer CropScience, and DFG for the development of iBeetle-Base (DFG LIS project 417202192).

## Conflict of interest statement

None declared.

## Acknowledgements

We thank Martin Milner for dsRNA injections.

## Notes

### Competing Interest Statement

The authors have declared no competing interest.

